# Changes to the microbiome of the stomach and intestine of Threespine Sticklebacks associated with infection by the cestode *Schistocephalus solidus*

**DOI:** 10.1101/290015

**Authors:** Megan A Hahn, Nolwenn M Dheilly

## Abstract

Helminth parasites are extremely diverse, prevalent, and infective of virtually all phyla. However, until recently, parasitic helminth infections have been studied as interactions between the host and the parasite, leaving out key players; microbes. Microbes play key roles in the biology of all living organisms, and this concept is challenging our understanding of host and parasite interactions. Because the microbiome controls many aspects of the physiology and behavior of their host, the discovery that many helminths can disrupt their host microbiome, either deliberately or as a side effect of infection, is of great significance. However, this phenomenon has been exclusively studied in helminth parasites that inhabit the gut, and hence share the same environment as their host microbiome. In this study we investigate the composition of the microbiome of threespine sticklebacks infected by the cestode *Schistocephalus solidus* who grows in the sterile body cavity of its intermediate host. Using a combination of qPCR and 16S sequencing approaches we show that *S. solidus* infection is associated with a significant increase in bacterial load and changes in diversity and composition of the microbiome of bpoth the stomach and intestine of sticklebacks.. We hypothesize that *S. solidus* interacts with host microbes either directly during the first 24 hours of infection, or indirectly, through modulation of the host immune system. The role of the stickleback microbiome in the outcome of *S. solidus* infection remains to be investigated.

**Author summary:** The classical paradigm assumes that symptoms associated with a parasite infection are due to the parasite itself. However, parasites can alter their host microbiome, meaning that microbes could be the ones responsible for associated diseases, and be the target for new therapeutic strategies. Here, we investigate the association between infection by *S. solidus* and the diversity and composition of the host gut microbiome. Our study reveals for the first time that a cestode parasite, that grows within the body cavity of its host, could be responsible for changes in the microbiome composition of its host. This suggests that the interplay between parasite and host associated microbes may be due to indirect mechanism, such as modulation of the host immune response. Functional studies are now necessary to determine the mechanisms behind parasite-associated alterations of their host microbiomes.

## Introduction

The growing recognition of the fundamental role of microbes in animals and plant biology is changing our understanding of life. The holobiont is defined as the combination of host, and all its associated microbes [1]. We are only starting to comprehend the potential of applying these concepts to host-parasite interactions to decipher the role of host-microbe, parasite-microbe, and host-parasite-microbe interactions in the ecology and evolution of parasites [2]. Microbes provide crucial services to the host, including disease resistance. For instance, there is considerable evidence of the defensive role of bacterial symbionts in insects [3-6]. We are now able to take advantage of this property and resistant mosquitoes carrying the endosymbiotic bacterium *Wolbachia* are being engineered and released in natural populations to reduce the spread of diseases such as malaria, dengue, and Zika [7-12]. In vertebrates too, there is growing evidence that the microbiome participates in resistance mechanisms against diseases. For example, the disruption of the host microbiome at early life-stages, or just before infection can increase the susceptibility to diseases [13-15] and transplantation of the microbiome of resistant individuals into susceptible individuals can be sufficient to provide protection [16].

Given the various direct and indirect roles of microbes in host defense, chances are that parasites have gained the ability to take advantage of microbes to increase their virulence. Indeed, evolutionary theories predict that parasite counter-defense strategies are shaped by the host defense mechanisms that they encounter at the time of infection, which explains the major role of immune evasion, and of manipulation of immune pathways in parasite virulence mechanisms [17]. Alteration of the host microbiome has been associated with parasitic helminth infection in various species including pigs, hamsters, cattle, goats, and sheep [18-24]. In rats, infection by the cestode *Hymenolepis diminuta* results in a shift from *Bacilli* to *Clostridia* species [25]. The liver fluke *Opisthorchis viverrini*, associated with hepatobiliary diseases and cholangiocarcinoma, modifies its host intestinal microbiome and promotes *Helicobacter pylori* infection in the liver [26-28]. Similar effects were observed when studying natural populations. For instance, infection by soil transmitted nematodes *Trichuris spp.* in human individuals in Malaysia is correlated with an increased microbial diversity in the gut and a differential abundance of a subset of OTUs, with enrichment in the family *Paraprevotellaceae* [24]. Kreisinger et al.[29] showed that alterations of the gut microbiome composition are species specific; wild mice (*Apodemus flavicollis*) naturally infected by three different helminths (the nematodes *Heligmosomoides polygyrus* and *Syphacia spp.*, and the cestode *Hymenolepis spp.*) had significantly different microbiomes. However, all these studies focused on parasites’ alteration of the microbiome of the tissue within which the parasite resides. Here, we characterized the stomach and intestine microbiomes of Threespine sticklebacks, *Gasterosteus aculeatus*, parasitized or not with the cestode parasite *Schistocephalus solidus*. Within 24 hours after ingestion of an infected copepod, *S. solidus* passes through the intestine wall and reaches the body cavity of the fish where it matures into an adult worm, infective for birds [30]. Thus, *S. Solidus* directly interacts with the gut microbiome of the threespine stickleback for a very brief period. In the body cavity *S. solidus* may, however, impact the host microbiome indirectly by modulating the expression of immune response genes [31-33].

## Results

Threespine sticklebacks were collected live from Cheney Lake in Anchorage Alaska and shipped back to Stony Brook University. Tissue collection was conducted after a three week acclimation period in order to allow *S. solidus* to transit into the body cavity and grow to a conspicuous size if the fish had only recently been infected. Threespine sticklebacks were then starved for 12 hours, euthanized in tricaine methane sulphonate solution (MS222) and dissected in sterile conditions. Stomach (St) and Intestine (In) collected from six non-infected fish (HGa) and six fish parasitized by *S. solidus* (PGa) were collected and processed for analysis of their microbiome composition. Swab samples of the body cavity of two non-infected and two parasitized individuals were also collected and used as negative controls.

### Real-time qPCR of 16S rRNA gene

We used a quantitative PCR (qPCR) approach with universal primers targeting the bacteria 16S rRNA gene as a proxy to investigate the relationship between bacteria abundance in the fish host and *S. solidus* infection, (Fig 1, S1 Fig). Indeed, variations in 16S abundance among samples may be due to changes in bacterial load or changes in bacterial community composition.

**Fig 1.**
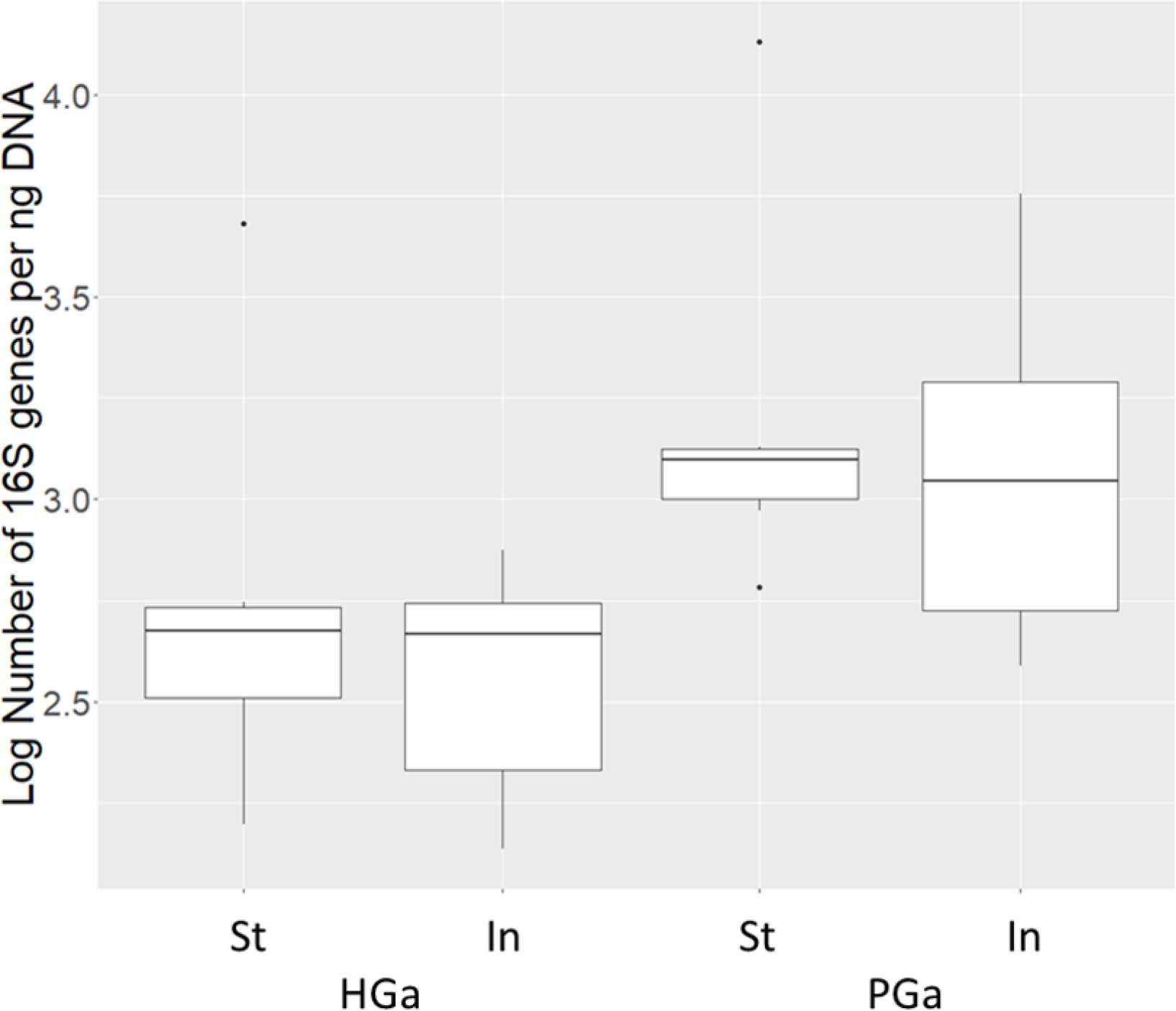
Bacterial load as estimated using qPCR of 16S gene. Bacterial load in the stomach (St) and intestine (In) of non-infected *G. aculeatus* (HGa), and parasitized *G. aculeatus* (PGa).

### Microbial diversity and composition

The bacterial community composition and diversity was determined by sequencing of the bacterial 16S rRNA gene V4 hypervariable region. Sequencing yielded an average of 43,645 (±SE 16,027) high quality paired-end reads for *G. aculeatus* tissue samples. Sequencing depth of control samples was higher and yielded an average of 83,875 (±SE 10,080) high quality reads. We resolved Amplicon Sequence Variants (ASVs) to avoid applying the arbitrary 97% threshold that defines molecular OTUs but has limited biological meaning [34]. The most abundant 100 ASVs in *G. aculeatus* represented a cumulative average of 85.1 % (±SE 1.1) of the paired-end reads in *G. aculeatus* tissue samples whereas it represented 7.8 % (±SE 0.4) reads in control samples. Conversely, the top 100 ASVs in control samples represented a cumulative average of 84.8% (±SE 1.4) of the paired-end reads in control samples, and only 4.4% (±SE 1.8) of the paired-end reads in *G. aculeatus* tissue samples. ASVs abundant in control samples were extremely similar to common bacterial contaminants of the MoBio kit used for bacterial DNA extraction and were removed from the analysis [35]. ASVs that were removed as contaminants can be found in S1 Table.

Alpha and beta diversity values were calculated on an ASV table rarified to 10,000 reads. Comparisons between non-infected and parasitized individuals yielded significant differences in alpha diversity values for Faith’s Phylogenetic Diversity (p<0.05, Table 1, Fig 2). Significant differences in Chao1 diversity were observed between stomach and intestine samples (p<0.05, Table 1, Fig 2).

**Fig 2.**
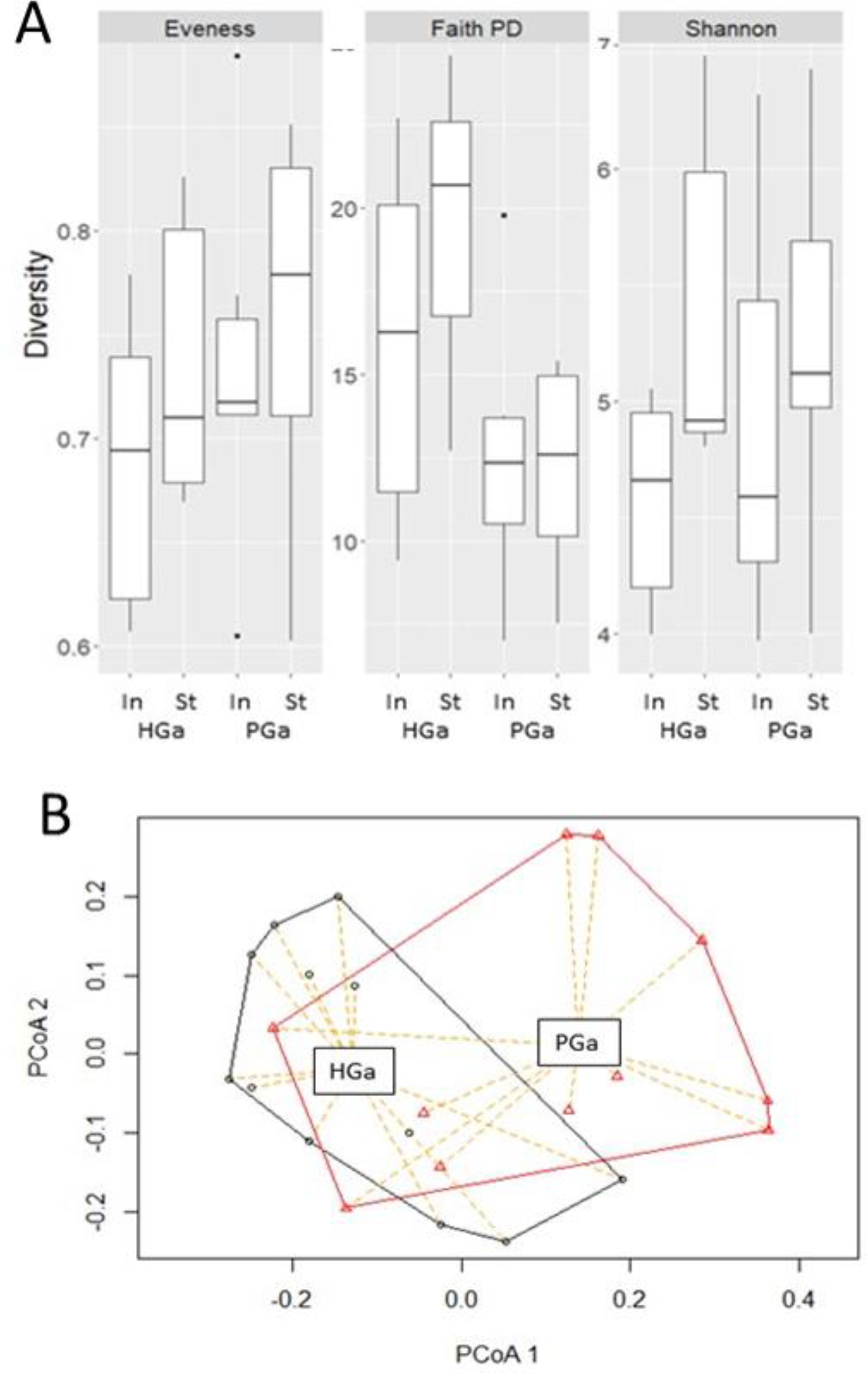
Comparison of alpha diversity metrics and principal coordinates analyses on beta-diversity metrics. A. alpha diversity values in the stomach (St) and intestine (In) of non-infected *G. aculeatus* (HGa), and parasitized *G. aculeatus* (PGa). B. Principal coordinates analysis based on Unweighted Unifrac distance at 10,000 rarefaction level showing differences between HGa and PGa.

**Table 1.** Results of two factor ANOVA on alpha diversity metrics of bacterial communities for all sample groupings including intestine (In), stomach (St), non-infected sticklebacks (H Ga), parasitized sticklebacks (P Ga) at the 10,000 rarefaction level.

| Comparisons |  | Evenness |  | Faith PD |  | Shannon |  | Chao1 |  |
| --- | --- | --- | --- | --- | --- | --- | --- | --- | --- |
| Group 1 | Group 2 | F | p-value | F | p-value | F | p-value | F | p-value |
| **H Ga** | **P Ga** | 1.040 | 0.319 | 8.562 | 0.008* | 0.086 | 0.772 | 1.199 | 0.287 |
| **Ga St** | **Ga In** | 1.152 | 0.295 | 0.753 | 0.395 | 3.470 | 0.077 | 6.614 | 0.018* |

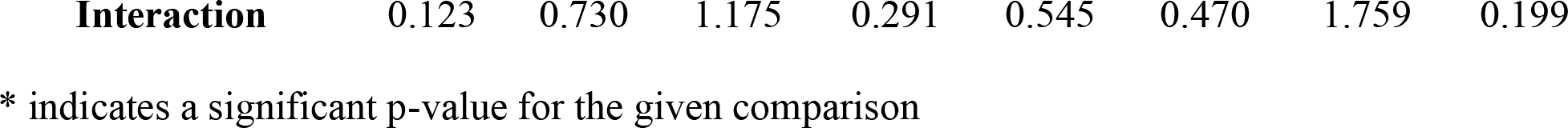

Principal coordinate analysis on weighted and unweighted UniFrac distances showed significant clustering of non-infected and parasitized tissue samples along PC1 (unpaired t-test p<0.05, Fig 2, S3 Fig and S3 and S4 Tables) and of intestine and stomach samples (Weighted UniFrac; PC2, paired t-test p<0.05, S5 and S6 Fig, S2 Table, Unweighted UniFrac; PC4 and PC17 paired t-test p<0.05, S3 Table). Community dissimilarity metrics (S5 Table) further confirmed that noninfected and parasitized individuals host significantly different microbiomes (PERMANOVA; weighted UniFrac P<0.05; unweighted UniFrac P<0.005) with significant differences in dispersion (PERMDISP; weighted UniFrac p<0.05, unweighted UniFrac p<0.05). These results indicate that the microbiome of parasitized individuals is overall less variable than the microbiome of noninfected individuals. Together with the reduced phylogenetic diversity, it indicates a significant reduction in abundance of variants of bacteria, rather than overall changes in species composition, No significant differences were observed between stomach and intestine samples using these metrics.

### Taxa distribution and differential abundance analysis

We used DESeq2 to identify bacteria phylotypes differentially abundant among samples. The proportion of ASVs differentially abundant between non-infected and parasitized *G. aculeatus*, was similar to the proportion of ASVs differentially abundant between intestines and stomachs of *G. aculeatus*, or differentially abundant between non-infected and parasitized individuals in stomach and intestine (Table 2, S6 and S7 tables).

**Table 2.** Number of differentially abundant ASVs/genus identified by DESeq2 (Adj p value <0.05).

| Comparison between |  | Total | More in A | More in B |
| --- | --- | --- | --- | --- |
| A | B |  |  |  |
| Healthy (H) | Parasitized (P) | 44 / 1 | 22 / 1 | 22 / 0 |
| Intestine (In) | Stomach (St) | 43 / 4 | 8 / 1 | 35 / 3 |
| Infection Status:Tissue Interaction |  | 48 / 3 |  |  |

Stickleback intestine and stomach microbiome communities were largely dominated by few ASVs of *Clostridiales, Rickettsiales, Burkholderiales* and *Desulfovibrionales* (Fig 3). The cumulative median relative abundance of the five most abundant ASVs in the threespine stickleback gut microbiome communities represented 56% of the reads, while the next five most abundant ASVs represented 20% of the reads. On few occasions, individual fish had high abundance of ASVs that only occurred in low abundance in other samples. For example, individual Ga12 (Fig 3) had high abundance of *Gammaproteobacteria* of the family *Shewanella* with a cumulative relative abundance in the stomach and intestine of 31.4% and 41.7%. Yet, the cumulative mean relative abundance across all other fish is 9.2% and 8.7% for the stomach and intestine, respectively.

**Fig 3.**
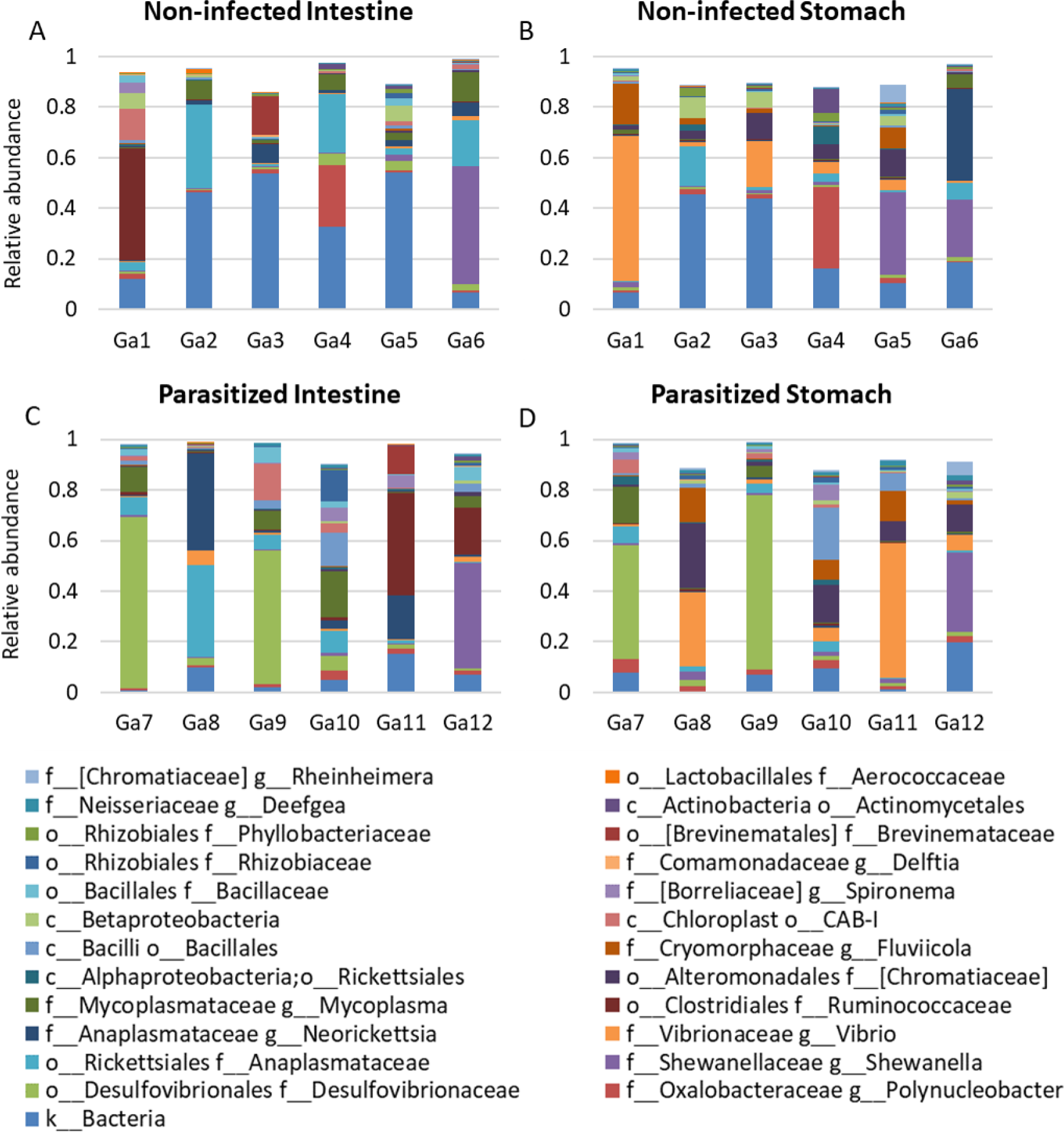
Relative abundance of the 25 most abundant ASVs in the intestine (A and C) and stomach (B and D) of non-infected (A and B, samples Ga1 to 6) and parasitized (C and D, samples Ga 7 to 12) Threespine Sticklebacks. The legend indicates the family (f_) and genus (g_) names.

Despite the very subtle differences in overall diversity between stomach and intestine samples, significant differences in relative abundance of 43 ASVs were observed, but only 3 genera of low abundance bacteria showed significant differences. Seventeen ASVs were found significantly more abundant in intestine than stomach and 26 ASVs were found significantly more abundant in stomach than intestine (Table 2). In particular, stomachs were enriched in various *Gammaproteobacteria* of the order *Alteromonadales*, in *Alphaproteobacteria* of the order *Rickettsiales* and *Sphingomonadales*, and a *Bacilli* of the order *Lactobacillales.* Intestines were enriched in *Clostridia*, of the order *Clostridiales*, and in *Cytophaga* of the order *cytophagale.* Of interest, different sequence variants of *Alphaproteobacteria* of the order *Rhizobiales, Betaproteobacteria* of the order *Burkholderiales*, and *Spartobacteria* of the order *Chtoniobacteriales* were enriched in the intestine and stomach of sticklebacks.

Similarly, only three low abundance bacteria were found differentially abundant between noninfected and parasitized individuals (Table 2, S6 table) suggesting that different variants of bacteria from the same genus are found in non-infected and parasitized individuals. Thirty-nine of the ASVs were less abundant in parasitized individuals, with a majority of *Alphaproteobacteria* of the order *Rickettsiales*, *Betaproteobacteria* of the order *Burkholderiales*, and *Gammaproteobacteria* of the orders *Alteromonadales* and *Oceanospirillales.* Twenty-two ASVs were more abundant in parasitized individuals including *Clostridia* of the order *Clostridiales*, and *Bacilli* of the order *Bacillales, Alphapoteobacteria* of the order *Rhizobiales* and *Sphingomonadales, Gammaproteobacteria* of the order *Aeromonadales* and *Spirochaetes* of the order *Borreliales*.

## Discussion

In this study, we analyzed the effect of infection by *S. solidus* on the composition of the stomach, (which is often omitted from gut microbial composition analyses), and intestine microbiome of its threespine stickleback host. Both RT-qPCR analyses used to quantify 16S rRNA gene abundance and 16S rRNA sequencing used to characterize microbiome diversity and composition revealed significant differences in the microbiome of both the stomach and intestine of the fish, even though the parasite was not in contact with these microbes anymore. Helminths have been found to impact the gut microbiome of humans and livestock [18-20, 22, 23, 26]. This modification of the host microbiome has been suggested to participate in strategies to impact the host immune system and establish infection [36]. However, these studies focused on intestinal parasites. In this study, we found that parasite infection is associated with an increase in abundance of 16S rRNA genes, but a reduction in phylogenetic diversity associated with a reduction of beta diversity and a small shift in bacterial microbiome composition. The correlation between our observation of changes in fish gut microbiome composition and infection status could result from two non-mutually exclusive hypotheses: non-infected individuals may carry defensive bacteria that reduce the chance of successful infection by *S. solidus*, or *S. solidus* infection may result in an alteration of the host microbiome. Our study identifies a number of ASVs significantly differentially abundant in the stomach and/or intestine of non-infected and parasitized individuals that could be targeted in future studies to test these hypotheses using laboratory controlled experimental infections.

Plerocercoids of *S. solidus* were not in contact with the host gut microbiome at the time of sample collection. Therefore, two hypotheses could be explored going forward to determine how *S. solidus* infection might impact its host microbiome. It is possible that *S. solidus* directly interacts with stomach and intestine microbes during the first 24 hours of infection, before it transits into the body cavity of the fish [30]. The host gut microbiome can have important effects on host fitness and health, including immune and metabolic function [37, 38]. Thus, early alteration of the host gut microbiome at the onset of infection could provide long-term advantages to the parasite. Alternatively, changes in the fish microbiome composition could result from cross-talks between the microbiome and the immune system. Indeed, *S. solidus* alters different pathways of its host immune system at different time points over the course of an infection [39], that could indirectly impact the fish ability to regulate its microbiome composition. Also, a comparison of germ-free and conventional threespine sticklebacks revealed variations in the recruitment of granulocytes, and in expression of immune response genes [40, 41](including IL-22) that play key roles in host protection and facilitation of helminth establishments[42, 43]. Further studies are therefore necessary to determine at which time point *S. solidus* infection results in changes in the stickleback microbiome.

It should be noted here that this study focused on field-caught individuals. Given the very high prevalence of *S. solidus* in Cheney lake [44], it is very likely that all non-infected individuals had been exposed to *S. solidus.* As such, parasite exposure, if not parasite infection, probably represents gut homeostasis in natural conditions. It means that more important differences can be expected from future laboratory control experiments focusing on the differences in the microbiome composition of healthy non-exposed sticklebacks, and both exposed but non-infected and successfully infected sticklebacks. Because high parasite prevalence is highly common in field samples of any species, and because this study, and many others have found a significant correlation between parasitism and changes in microbiome composition, we would like to conclude by emphasizing that the presence of macro-parasites in field caught individuals should always be assessed before conducting a microbiome study in natural conditions.

## Methods

### Sample collection

In September 2015, threespine stickleback (*Gasterosteus aculeatus*) specimens were collected live from Cheney Lake in Anchorage Alaska (61° 12’ 17” N, 149° 45’ 33” W). Fish were caught using un-baited minnow traps placed along the north shoreline of the lake between 0.25 and 2 m deep, and at least 2 m apart. As stickleback have relatively short lifespans and it is easy to distinguish young of the year fish from fish that are 1+ or 2+ years old, we took fish that ranged from 3-4 cm in length to ensure that we sampled from the same cohort [45]. Fish were shipped to Stony Brook University, kept in 20-gallon tanks of 6% seawater at 5°C at the Flax Pond Marine Lab, and fed mysis shrimp twice per day.

Following an acclimation period of 3 weeks, and 12 hours of starvation, fish were anaesthetized and euthanized in tricaine methane sulphonate solution (MS222), decapitated, and dissected in sterile conditions. Prior to dissections all surfaces were cleaned with 80% ethanol. Between each subsequent fish dissection, tools and dissecting plate were cleaned with 80% ethanol and betadine to remove potential contaminants. Before dissection, fish were brushed with betadine to prevent contamination of the body cavity with microbes found on the surface of the fish. For dissection, an incision was made along the lateral line of the fish body, around the bony pelvis. The cut extended from the pectoral fins to just anterior of the anus to avoid cutting the intestine. After the initial incision, the bony pelvis was pulled away, opening up the body cavity. The sex of the fish was assessed by visual inspection of gonads at the time of dissection and then confirmed using PCR with sex specific primers as described in (REF). The presence of *S. solidus* was recorded, and stomach and intestines were flash frozen in liquid nitrogen for future DNA extraction. The number of parasites per infected fish is provided in table 3. Swab samples of the body cavity of two noninfected and two parasitized individuals were also collected. All samples were then stored at -80°C until use.

**Table 3.** Sex of threespine sticklebacks used in this study and number of *S. solidus* parasite found within parasitized individuals.

| Non-infected Fish | Sex | Parasitized Fish | Sex | # of parasites |
| --- | --- | --- | --- | --- |
| Ga1 |  | Ga7 |  | 3 |
| Ga2 |  | Ga8 |  | 4 |
| Ga3 |  | Ga9 |  | 4 |
| Ga4 |  | Ga10 |  | 1 |
| Ga5 |  | Ga11 |  | 1 |
| Ga6 |  | Ga12 |  | 2 |

\* M: Male; F: Female

### Ethics statement

Stickleback collection followed guidelines for scientific fish collection by the State of Alaska Department of Fish and Game in accordance with Fish sampling permit #SF2015-263 and #P17-025 and fish transport permits #15A-0084 and 17A-0024 provided to NMD. Fish were maintained at Stony Brook University under the License to collect or possess #1949 provided by the New York State Department of Environmental Conservation to NMD. Fish experiments were conduct following protocols described in Institutional Animal Care and Use Committee (IACUC) #237429 and # 815164 to Michael Bell and NMD, respectively. Fish euthanasia was conducted using MS222 and decapitation before parasite and tissue sampling. All experiments were performed in accordance with relevant guidelines and regulations in the Public Health Service Policy (PHS) on Humane Care and Use of Laboratory Animals.

### DNA extraction

Tissues from six non-infected fish (HGa) and six infected fish (PGa) were collected for DNA extraction. DNA purification from stomach (St) and intestine (In) tissues was performed using a MoBIO Powerlyzer Powersoil Kit (Cat#12855) with the following modifications to the manufacturer’s protocol. First, all tissue samples were placed in tough tubes with 3mm glass beads. To each tube, we added 750μL of the bead beating solution supplied with the extraction kit. Samples underwent three rounds of 20 sec bead beating cycles and were kept on ice between each round. The homogenate was transferred to the MoBIO supplied 0.1mm bead tube and incubated with 60μL of the provided C1 solution for 10 min at 60°C. After vortexing, centrifugation, and filtration with solution C5 according to manufacturer’s protocol, 75 μL of solution C6 was added to the spin filter in a clean 2mL tube and incubated at room temperature for 5 min before centrifugation at 10,000g for 30 sec. This final step was repeated to obtain a final DNA extract of 150μL.

### Real-time qPCR

Bacteria from all samples were quantified by qPCR using the QuantStudio 6 Flex RT PCR System (Fisher Scientific). Two 16S primers were used (Forward: TCCTACGGGAGGCAGCAGT, Reverse: GACTACCAGGGTATCTAATCCTGTT [46] and Forward: GTGSTGCAYGGYTGTCGTCA, Reverse: ACGTCRTCCMCACCTTCCTC [47]) and compared to an *E. coli* standard for which cell count was obtained using a hemocytometer (Stony Brook, NY). The qPCR mixture (20μL) was composed of 10μL of SYBR green master mix (CAT# 4309155), 4.8μL Molecular biology grade water (CAT#46000), 0.6μL forward primer, 0.6μL reverse primer, and 4μL DNA.

### 16S rDNA amplification and Illumina MiSeq sequencing

We sequenced the stomach and intestine microbiota from six non-infected and six parasitized individuals. Swabs of the body cavity, and negative controls from the DNA extraction were also sequenced. The 16S rDNA V4-V5 hypervariable region was amplified with *E. coli* 515f and 806r primers (with barcode on the forward primer) as specified by the Earth Microbiome Project [48-51]. We used a 28 cycle PCR (5 cycle used on PCR products) using the HotStarTaq Plus Master Mix Kit (Qiagen, USA) under the following conditions: 94°C for 3 minutes, followed by 28 cycles of 94°C for 30 seconds, 53°C for 40 seconds and 72°C for 1 minute, after which a final elongation step at 72°C for 5 minutes was performed. After amplification, PCR products are checked in 2% agarose gel to determine the success of amplification and the relative intensity of bands. Multiple samples are pooled together (e.g., 100 samples) in equal proportions based on their molecular weight and DNA concentrations. Pooled samples are purified using calibrated Ampure XP beads. Then the pooled and purified PCR product is used to prepare illumina DNA library. Sequencing was performed at MR DNA to obtain 250bp paired end reads (www.mrdnalab.com, Shallowater, TX, USA) on a MiSeq following the manufacturer’s guidelines. Sequence data were initially processed using MR DNA analysis pipeline (MR DNA, Shallowater, TX, USA). In summary, sequences were joined, depleted of barcodes then sequences <150bp removed, sequences with ambiguous base calls removed.

### Microbiome composition analysis

The 16S rDNA gene Illumina reads were further processed using methods implemented in QIIME2, Phyloseq [52, 53] and R as detailed in the supplementary information (Supp. File 1). Briefly, samples were demultiplexed and quality filtered. ASVs (amplicon sequence variants) were picked against the Greengene database using the DADA2 pipeline.. The final ASV table was then rarefied to a depth of 3000 sequences per sample for comparisons between the microbiomes of *S. solidus* and stickleback tissue samples, and 10,000 sequences for comparisons among stickleback tissue samples. Rarefaction curves showed that at these depths of sampling, we were able to sample a large portion of the ASV diversity present while still retaining all samples (Supp File 2).

### Diversity measures and statistical analyses

Analyses were performed using QIIME2 and the Phyloseq and DESeq2 packages in R [53, 54]. Each of the following diversity metrics and statistical analyses were carried out at the 10,000 rarefaction levels to perform comparisons (i) among all sample categories (HGaIn, PGaIn, HGaSt, PGaSt), (ii) between tissue (all In vs all St), and (iii) depending on infection status (all HGa versus all PGa). Pielou’s Evenness, Shannon, Simpson, and Faith’s Phylogenic Diversity (PD_Whole_tree, with Greegene taxonomy tree) alpha diversity metrics were used to estimate within species diversity. Alpha diversity was compared between sample types using pairwise Kruskal-Wallis H-tests. Beta diversity was calculated with both weighted and unweighted UniFrac metrics. PERMANOVA at 10,000 permutations was carried out to identify significant differences between categories of samples. We tested for homogeneity of multivariate dispersion using PERMDISP. Principal coordinate analysis (PCoA) was performed to look for patterns in unconstrained multivariate space and a combination of paired and unpaired pairwise t-tests were used to test if axes significantly discriminate groups of samples. DESeq2 was used on raw data to examine differences in individual microbe abundance.

## Data Accessibility

All data is being made available through the NCBI database under BioSample accessions BioSample accessions SAMN08800337 to SAMN08800360, SAMN08805227, SAMN08805228, SAMN08805230 and SAMN08805232.

## Acknowledgements

We would like to acknowledge the expertise and assistance of Dr. Michael Bell in fish collection and dissection techniques.

## Supporting information

**S1 File: Document contains S1 to S7 Figs with corresponding legends**

**S2 File: Contains S1 to S9 tables with corresponding legends**

